# Transposons repressed by H3K27me3 were co-opted as cis-regulatory elements of H3K27me3 controlled protein coding genes during evolution of plants

**DOI:** 10.1101/2022.10.24.513474

**Authors:** Tetsuya Hisanaga, Facundo Romani, Shuangyang Wu, Teresa Kowar, Ruth Lintermann, Bhagyshree Jamge, Sean A. Montgomery, Elin Axelsson, Tom Dierschke, John L. Bowman, Takayuki Fujiwara, Shunsuke Hirooka, Shin-ya Miyagishima, Liam Dolan, Daniel Schubert, Frédéric Berger

## Abstract

The mobility of transposable elements (TEs) contributes to evolution of genomes ^1,2^. Meanwhile, their uncontrolled activity causes genomic instability and therefore expression of TEs is silenced by host genomes ^3,4^. TEs are marked with DNA and H3K9 methylation that are associated with silencing in flowering plants ^5^, animals, and fungi ^6^. Yet, in distantly related eukaryotes TEs are instead marked by H3K27me3 deposited by the Polycomb Repressive Complex 2 (PRC2) ^7–11^, an epigenetic mark associated with gene silencing in multicellular eukaryotes ^12–15^. It was therefore proposed that the ancestral activity of PRC2 was the deposition of H3K27me3 to silence TEs ^16^.

To test this hypothesis we obtained mutants deprived of PRC2 activity and used genomics to analyze the role of PRC2 in extant species along the lineage of Archaeplastida. While in the red alga *Cyanidioschyzon merolae* more TEs than genes were repressed by PRC2, an opposite trend was observed in bryophytes *Marchantia polymorpha* and *Anthoceros agrestis*. In the red alga, TEs silenced by H3K27me3 are in subtelomeres but in bryophytes, TEs and genes marked by H3K27me3 form coregulated transcriptional units. The latter trend was also observed in the flowering plant *Arabidopsis thaliana*, and we identified cis-elements recognised by transcription factors in TEs flanking genes repressed by PRC2.

Together with the silencing of TEs by PRC2 in ciliates that diverged early from an ancestor common with Archaeplastida, our findings support the hypothesis that PRC2 deposited H3K27me3 to silence TEs in early lineages of eukaryotes. During evolution, TE fragments marked with H3K27me3 were selected to shape transcriptional regulation that control networks of genes regulated by PRC2.

**Highlights:** H3K27me3 marks a decreasing proportion of TEs during evolution of plants

The polycomb repressive complex 2 represses TEs in red algae and bryophytes

H3K27me3-marked TEs in flowering plants contain transcription factor binding sites

Transcription factors bind TEs and regulate networks of genes controlled by PRC2

## Results and Discussion

### PRC2 represses expression of TEs in the red alga *Cyanidioschyzon merolae*

To investigate the evolution of PRC2 function in the Archaeplastida lineage, we first selected the red alga *Cyanidioschyzon merolae*, belonging to the class Cyanidiophyceae, that diverged from the Viridiplantae ca. 1.200 MYA ^17^. Previous genome-wide profiling of H3K27me3 demonstrated that this mark is associated not only with Protein Coding Genes (PCGs) but also with transposable elements (TEs) and repeats ^18^. To re-analyse these data we updated the annotation of TEs in the genome of *C. merolae* (see details in Methods). All 1755 TEs annotated belonged to class II DNA transposons with 34% of TEs from the Mutator family, 24% helitrons, 2.5% CACTA and 40% are unknown (Figure S1A). We observed a broad overlap between TE density and H3K27me3 coverage over whole chromosomes (Figure 1A). The overlap between TE or PCG annotations and H3K27me3 peaks showed 4.7% of PCGs and 48% of TEs were covered by H3K27me3 and there was no significant enrichment of specific TE families among them (Figure S1A). Because half of the TEs were not covered by H3K27me3 we investigated the presence of other repressive marks such as methylation of the lysine 9 of histone H3 (H3K9me1) and DNA (5methyl-C (5mC)), which mark TEs in many eukaryotes including flowering plants ^5,6^. In the genome of *C. merolae*, five putative SET domain histone methyltransferases are encoded (four with homology to H3K4 methyltransferases, one with homology to the PRC2 methyltransferase Enhancer of Zeste (E(z)). Thus, none has homology to KRYPTONITE/SU(VAR)3-9 HOMOLOG 4, which is responsible for H3K9 methylation in *A. thaliana*. Accordingly, we detected higher levels of H3K27me3 and H3K27me1 than H3K9me1 via immunoblotting (Figure 1B). Furthermore, we re-analyzed a genome-wide 5mC profile of *C. merolae* ^*19*^, and confirmed that 5mC levels of the nuclear genome are not higher than the background levels measured on chloroplast and mitochondria DNA (Figure S1B). We conclude that it is unlikely that DNA methylation has a role in TE silencing and, furthermore, H3K27me3 is more broadly associated with TEs than with genes in *C. merolae*.

**Figure 1.**
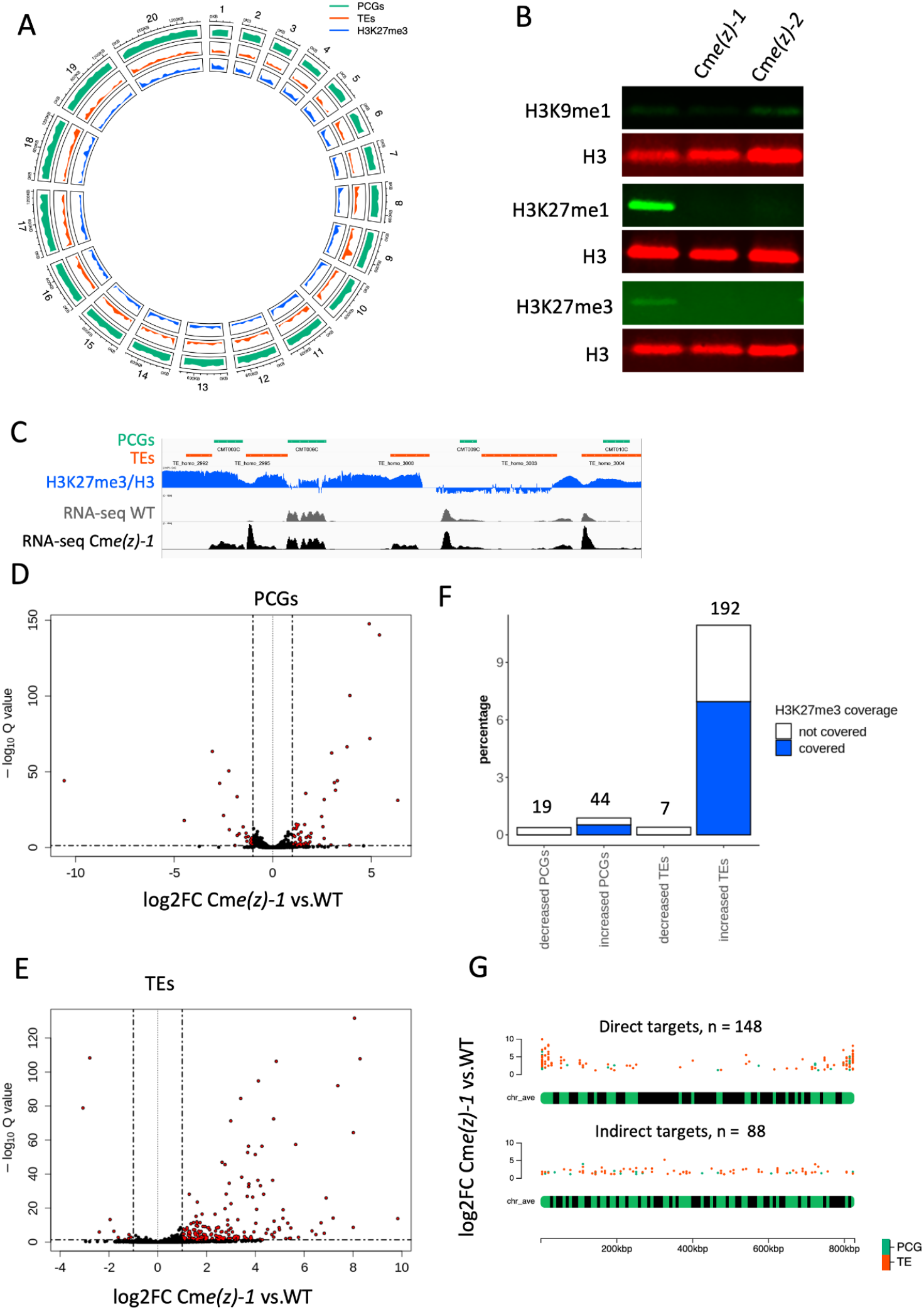
PRC2 represses TE expression in the red alga. (A) Circos plot showing genome wide distributions of protein coding genes (PCGs, green), transposable elements (TEs, pink) and H3K27me3 peaks (blue). Each band shows the density of annotated PCGs, TEs or chromatin mark peaks per chromosome, relative to the greatest density per band. (B) Protein gel blot analyses indicating level of each histone modification in the wild type and two independent loss-of-function Cm*e(z)* mutants. (C) A screenshot of integrative genomics viewer showing subtelomeric region of chr 20. The tracks show TE annotation (orange), gene annotation (green), H3K27me3 coverage (blue), expression level in wild type and Cm*e(z)-1* mutant (grey), respectively. H3K27me3 coverage is normalized against the H3 ChIP signal. (D) Volcano plot showing differential expression of protein coding genes (PCGs) between wild type and Cm*e(z)-1* mutant. Differentially expressed PCGs (q. value < 0.05, and |log2 fold change| > 1) are marked in red. (E) Volcano plot showing differential expression of transposable elements (TEs) between wild type and Cm*e(z)-1* mutant. Differentially expressed TEs (q. value < 0.05, and |log2 fold change| > 1) are marked in red. (F) A bar plot indicating percentage of PCGs and TEs exhibiting decreased or increased expression levels in Cm*e(z)-1* mutant in all PCGs and TEs. Those who are not covered or covered by H3K27me3 are shown in white or blue, respectively. (G) A chromosomal plot showing relative positions of PRC2 direct targets (top) and indirect targets (bottom). X-axis indicates relative positions of each target on an artificial chromosome which has averaged size. Y-axis indicates log2-fold change in RNA-seq analysis comparing Cm*e(z)-1* vs. wild type.

To explore the role of H3K27me3 in transcriptional repression of TEs in *C. merolae*, we disrupted the only ortholog of E(z), Cm*E(z)* (CMQ156C) (Figures S1C and S1D). Two independent loss of function alleles Cm*e(z)-1* and Cm*e(z)-2* exhibited a near complete loss of H3K27me3 with a concomitant reduction of H3K27me1 levels, but no decreased levels of H3K9me1 (Figure 1B), suggesting that Cm*E(z)* deposits H3K27me3 but not H3K9me1. The low levels of H3K27me1 in the mutant are likely explained by demethylation of H3K27me3 by JUMONJI demethylases that remain to be characterized in *C. merolae*. A comparison of transcriptomes of Cm*e(z)-1* and wild type showed that expression levels of 0.4% (19) and 0.8% (44) protein coding genes (PCGs) out of a total 4984 PCGs decreased and increased, respectively (Q.value < 0.05, |log2 fold change| > 1; Figures 1C - 1E). None of 19 PCGs with decreased expression in Cm*e(z)-1* and about 60% (26) of PCGs exhibiting increased expression in Cm*e(z)-1* were covered by H3K27me3 in wild type (Figure 1F), indicating that PRC2 represses expression of a small number of PCGs via deposition of H3K27me3. The loss of Cm*E(z)* also increased expression of 192 out of 1755 TEs, while it resulted in decreased expression of only 7 TEs (Figures 1E and 1F). More than 60% (122) of all TEs showing increased expression in Cm*e(z)-1* were covered by H3K27me3 in the wild type (Figure 1F), supporting that PRC2 represses the expression of TEs. We defined PCGs and TEs, which exhibited both increased expression in Cm*e(z)* and were covered by H3K27me3 in the wild type as direct targets of PRC2 (26 PCGs and 122 TEs). We observed no enrichment of these PRC2 direct targets in a specific TE family (Figure S1A). Most TEs repressed by PRC2 were located in subtelomeric chromosomal regions (Figure 1G). By contrast, indirect targets of PRC2 that were misexpressed were scattered along chromosomes (Figure 1G). We concluded that in *C. merolae* PRC2 deposits H3K27me3 and represses expression of TEs preferentially in subtelomeric regions.

### Association of H3K27me3 marks on TEs are conserved among bryophytes

In bryophytes, that diverged from vascular plants ca. 500-460 MYA and comprise hornworts, liverworts and mosses ^20^, a large fraction of TEs are covered by H3K27me3 in the liverwort *Marchantia polymorpha* ^9^ while TEs in the model moss *Physcomitrium patens* are covered mostly by H3K9me2 ^21^. To address the conservation of the association of H3K27me3 with TEs among bryophytes, we used the model hornwort *Anthoceros agrestis* that diverged from bryophyte ancestors before the divergence of liverworts from mosses ^22^. We annotated TEs in *A. agrestis* Oxford strain and identified 90,809 TEs including 990 intact TEs belonging to various TE families (Figure S1E). By contrast with *C. merolae*, we observed a much larger diversity of TE families and half of intact TE were retrotransposons. This enrichment of LTR TE families was also observed in TEs longer than 500 bp (total 26,070 TEs including 923 intact TEs) (Figure S1E) and thus we analyzed separately longer TEs and shorter TEs. Using chromatin immunoprecipitation coupled with DNA sequencing (ChIP-seq), we obtained genomic profiles of five histone post translational modifications (PTMs) (H3K4me3, H3K36me3, H3K9me1, H3K27me1 and H3K27me3) and H3 from four week old vegetative tissue of *A. agrestis* (see ^23^ for a general overview of the chromatin of *A. agrestis*). We performed k-means clustering of chromatin marks over TEs, and defined five major clusters of TEs showing different chromatin environments (Figure 2A). Clusters 2 and 3 contained 39% and 25% of all TEs and were covered with H3K9me1 and H3K27me1 (Figure 2A). Cluster 1 comprised 3% of all TEs that were not covered with H3K9me1, H3K27me1 and H3K36me3, but with H3K4me3 and H3K27me3. We did not find a preferential association between TEs from cluster 1 and specific TE families. More than 80% of TEs in cluster 1 are located close to genes belonging to PCG cluster 1 that are also covered with H3K27me3 (Figure 2C). According to their chromatin environment, short TEs clustered in a fashion similar to long TEs (Figure S2B) and the short TEs covered with H3K27me3 and H3K4me3 were also associated with PCGs sharing the same chromatin environment (Figure S2B). Similarly, we observed the association of 5395 TEs with PCGs covered by a chromatin landscape dominated by H3K27me3 in *M. polymorpha* (TE Cluster 1 and PCG Cluster 1 Figure 2B, S2B ^9^). We concluded that the association of H3K27me3 with TEs is conserved in liverworts and hornworts. Because hornworts are sister to the other two groups ^22^, the association of TEs with H3K27me3 would be ancestral to the bryophyte lineage.

**Figure 2.**
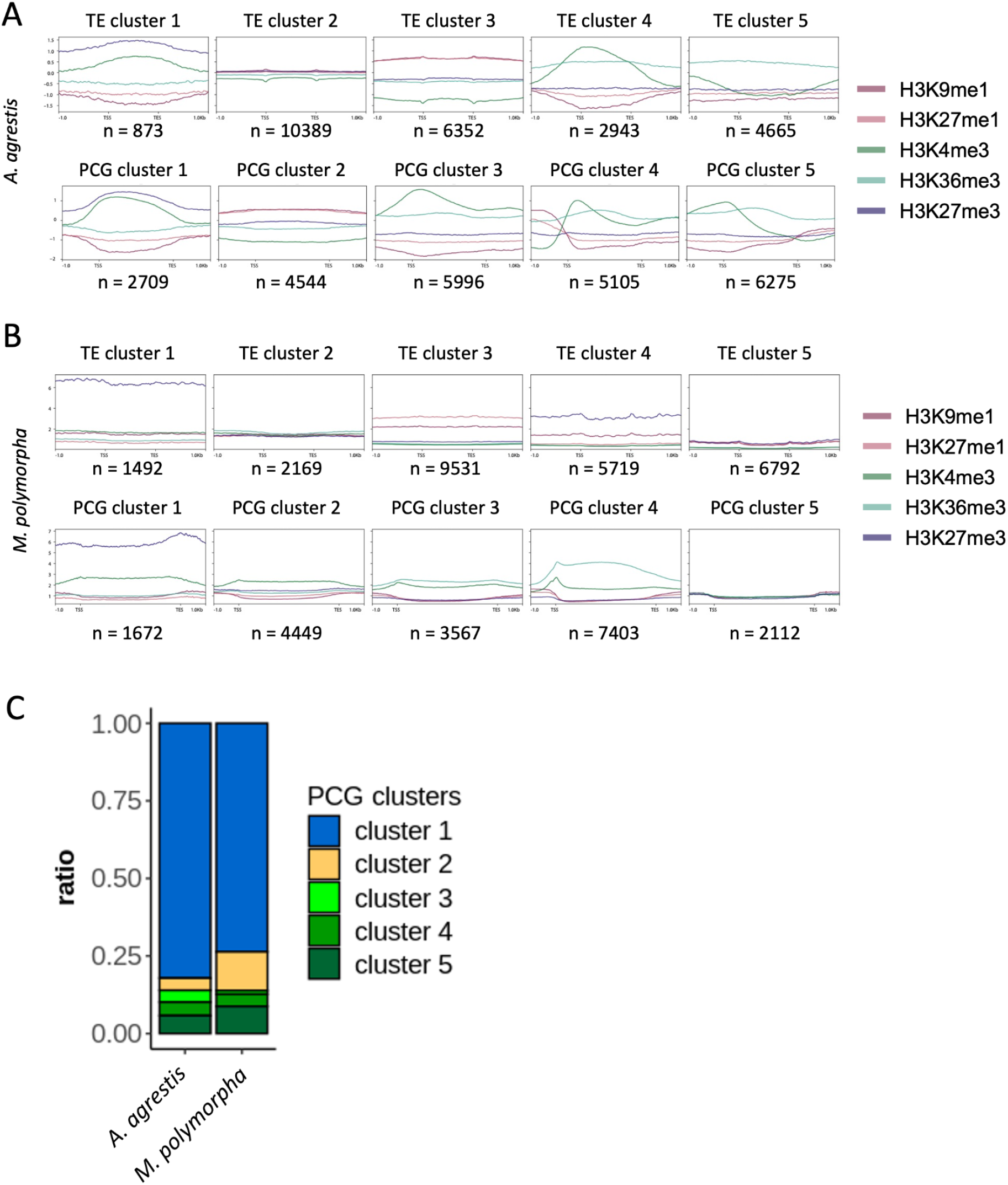
Chromatin landscape of Bryophytes. (A) Aggregate profile plots showing log2 ChIP/H3 enrichment for various chromatin modifications per TE (top) or PCG (bottom) cluster in *A. agrestis*. (B) Aggregate profile plots showing log2 ChIP/H3 enrichment for various chromatin modifications per TE (top) or PCG (bottom) cluster in *M. polymorpha*. (C) Stacked bar chart showing proportion of gene clusters of nearby genes of TEs in TE cluster 1 in *A. agrestis* (left) and *M. polymorpha* (right).

### PRC2 represses expression of TEs in *M. polymorpha*

To test whether PRC2 silences TEs in bryophytes, we used *M. polymorpha* which is amenable to genetic manipulation ^24^. To disrupt the function of PRC2, we focused on the ortholog of the PRC2 catalytic subunit Enhancer of zeste, E(z) that is encoded by three *E(z)* paralogs in *M. polymorpha* genome ^25^. Only Mp*E(z)1* is expressed in vegetative gametophytic tissues of *M. polymorpha ^26^*. A knock-down of Mp*E(z)1* showed reduced growth and necrosis of tissues that hindered further analysis ^27^. We observed that genes encoding the homeodomain transcription factors KNOX and BELL which are not expressed in the vegetative tissues of the gametophyte were covered by H3K27me3 (Figure S3A). Since these transcription factors are essential in determining life phase transitions in plants ^28–31^, we hypothesized that their mis-expression in Mp*e(z)1* mutant could be responsible for the lethality observed in the knock down of Mp*E(z)1*. Therefore, we disrupted Mp*KNOX2* (see STAR methods for details) to obtain null Mp*knox2-1* alleles in Tak-2 female wild-type strain. In this mutant background we generated two knockout alleles of Mp*e(z)1*, Mp*knox2-1* Mp*e(z)1-1* and Mp*knox2-1* Mp*e(z)1-2* (Figures S3B and C). We also used a construct expressing two guide RNAs targeting Mp*KNOX2* and Mp*E(z)1* to obtain the combination of two additional alleles Mp*knox2-2* Mp*e(z)1-3* in the Tak-1 male wild-type strain (Figures S3B and S3C). While Mp*knox2* single mutants exhibited no developmental defects during the vegetative growth phase, Mp*knox2* Mp*e(z)1* double mutants exhibited slower thallus growth but with a morphology similar to the wild type (Figures S3D and S3E). This supported that misexpression of mpKNOX2 was responsible for the lethality observed in the Mp*e(z)1* null mutant.

Protein gel blot analyses using isolated nuclei from 14-day-old thalli indicated that compared with the wild type, H3K27me3 was undetectable in Mp*knox2-1* Mp*e(z)1*, while the levels of H3K9me1 and H3K27me1 were not reduced in Mp*knox2-1* Mp*e(z)1* (Figure 3A). In addition the levels of the three post transcriptional modifications analyzed did not change in wild type and Mp*knox2* (Figure 3A), indicating a specific and likely complete loss of H3K27me3 in the Mp*e(z)1* knockout mutant.

**Figure 3.**
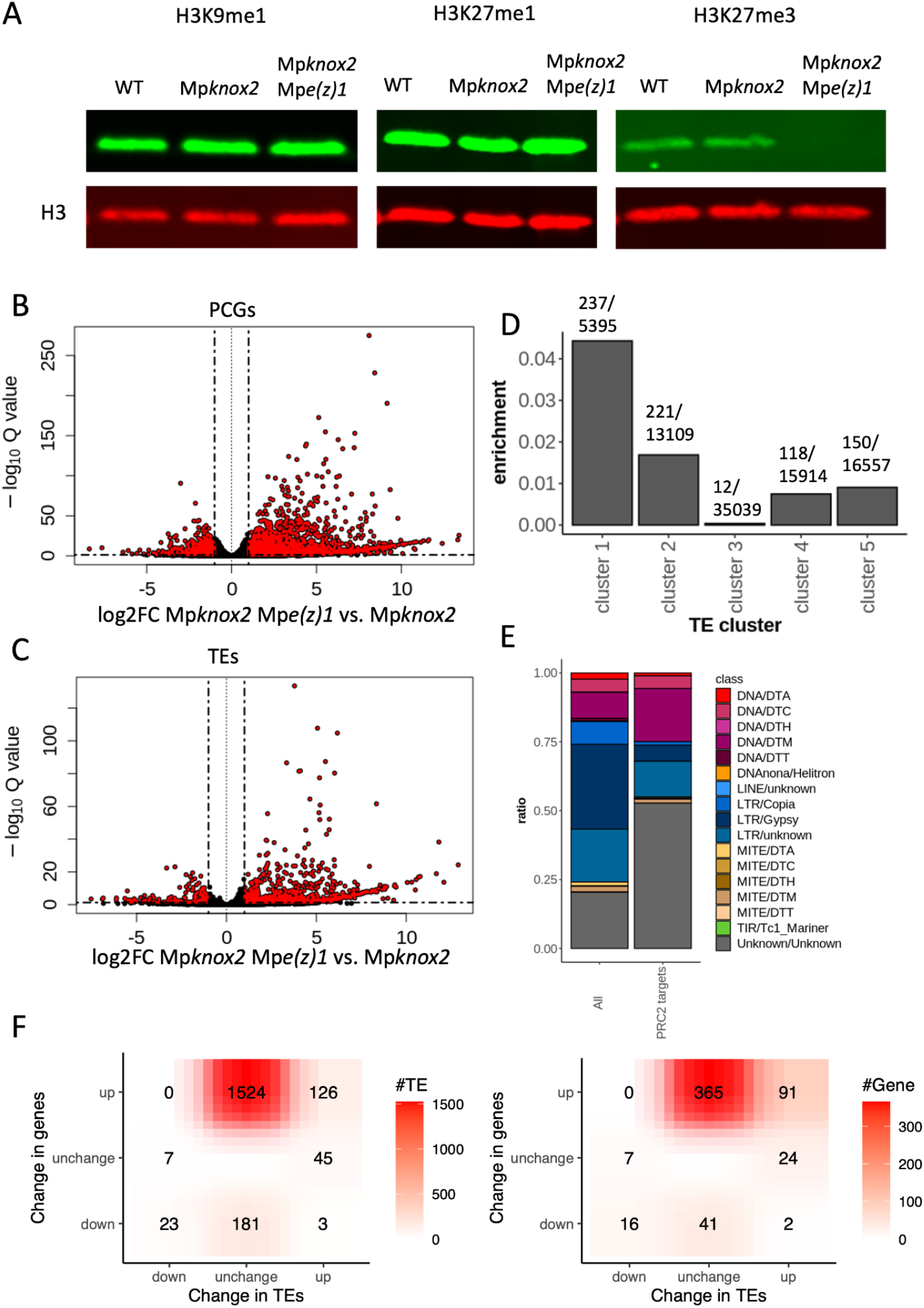
PRC2 silences TEs in *M. polymorpha*. (A) Protein gel blot analyses indicating level of each histone modification in the wild type, Mp*knox2* and Mp*knox2* Mp*e(z)1* mutant. (B) Volcano plot of all protein coding genes (PCGs) differentially expressed in Mp*knox2* Mp*e(z)1* compared to Mp*knox2*. Differentially expressed PCGs (Q. value < 0.05 and |log2FC|>1) are marked in red. (C) Volcano plot of transposable elements (TEs) differentially expressed in Mp*knox2* Mp*e(z)1* compared to Mp*knox2*. Differentially expressed (Q. value < 0.05 and |log2FC|>1) TEs are marked in red. (D) Bar plot indicating enrichment of up-regulated TEs in each TE cluster, calculated as the number of TEs exhibiting increased expression in each cluster divided by the total number of TEs in each cluster. (E) Stacked bar chart indicating proportion of TE families in all annotated TEs (All) and TEs repressed by PRC2 (PRC2 targets). Total numbers of TEs in each category are shown on bars. (F) The differential expression statistics of TE and neighbor PCGs (Left is TE number, right is the PCG number accordingly.) with hypergeometric test. P-value of the upregulated TE and neighbor PCGs pair is 0.

To evaluate the impact of the loss of H3K27me3 marks on the expression of PCGs and TEs, we compared the transcriptomes of the Mp*knox2* single mutant and the Mp*knox2-1* Mp*e(z)1* double mutant using total RNA isolated from 14 day old vegetative tissue. In the Mp*knox2* Mp*e(z)1* double mutant, 2240 PCGs showed increased expression compared to Mp*knox2* single mutant (Figure 3B) and 27% (606) of these genes were covered by H3K27me3 in wild-type thallus (Figure S3F), supporting the hypothesis that PRC2 represses transcription of PCGs via deposition of H3K27me3. PCGs repressed by PRC2 encoded proteins involved in secondary metabolism and response to various stresses (Figure S3G), suggesting a role of PRC2 in the response of vegetative tissues to the environment. We also observed the impact of the loss of PRC2 on the expression of TEs, using a new set of annotated TEs in *M. polymorpha* (see details in the methods). Expression of 738 TEs increased in the Mp*knox2* Mp*e(z)1* mutant compared with the Mp*knox2* mutant (Q.value<0.05, log2FC>1; Figure 3C). We confirmed the increased expression of several TE fragments and intact TEs using quantitative real time RT-PCR analysis (Figure S3H). We observed that TEs exhibiting increased expression in the Mp*e(z)1* mutant were primarily marked by H3K27me3 (TE Cluster 1, Figure 3D). These TEs belonged to mutator DNA transposons and other uncategorised TE families (Figure 3E and Fischer’s test). Because TEs and PCGs are interspersed in the genome of Marchantia, we investigated whether PRC2 coregulated TEs with their closest neighbor PCGs. We observed that three quarters of TEs from the TE cluster 1 were located close to genes from the PCGs cluster 1 (Figures 2E and S2B). Importantly, there was a significant enrichment of pairs of up-regulated TEs and neighbor PCGs (Figure 3F). We conclude that in *M. polymorpha*, PRC2 represses transcription from TEs. These are usually TEs associated with a PCG that is also repressed by PRC2.

### In *A. thaliana* TEs covered by H3K27me3 contain transcription factors binding sites

In flowering plants TEs are primarily marked by H3K9me1/2 but in mutants devoid of DNA methylation H2K27me3 becomes associated to TEs, suggesting that a link between TEs and PRC2 is masked by the presence of marks of constitutive heterochromatin. Based on a comprehensive analysis of chromatin states in *A. thaliana* we established that 11% of TEs are covered by H3K27me3 mark (Figure S4A and S4B, Cluster 6; and clustering details in ^32^). These TEs belong primarily to DNA and rolling circle (RC) transposon families in contrast to TEs from other clusters 1 - 3 that are covered with H3K9me1 and H3K27me1 (Figures S4C and S4D). Most of the TEs from cluster 6 were located nearby PCGs which are also covered by H3K27me3 (Figure S4E), suggesting that those TEs would have been co-opted to *cis-*regulatory motifs for nearby PCGs. To test whether these TEs could function as *cis*-regulatory elements controlling the repression of contiguous PCGs, we searched for binding of transcription factors in all TEs in *A. thaliana* and calculated enrichment of TF binding from publicly available ChIP-seq experiments in each TE cluster (Figure 4A). We observed a higher occupancy of TF binding in the TEs from cluster 6, which are marked with H3K27me3, than other TEs clusters (Figure 4A). This suggested H3K27me3-marked TEs function as *cis*-regulatory elements. Moreover, this bias was even clearer when we considered the functional TF binding events associated with the co-expression of the neighboring gene (Figure 4A). By contrast TF binding events were not found in TEs from clusters 1 - 4, which are covered with constitutive heterochromatin (H3K9me1/2 and H3K27me1). Altogether these observations suggested that TFs associated with TEs are important for the regulation of neighboring genes. Among TFs identified enriched in TEs from cluster 6, we found four MADS-box containing TFs (APETALA1, SEPALLATA3, AGAMOUS-LIKE15, and PISTILLATA), which are known to interact with PRC2 and control flower development (Figure 4B) ^33,34^. A high proportion of TEs marked with H3K27me3 was located near PRC2 regulated PCGs covered by the Chromatin Landscape 1, defined by enrichment in H2A.Z, H3K27me3 and H2AK121Ub, and excluded from actively transcribed genes associated with chromatin landscapes 7 - 10 (defined in ^32^) (Figures 4C and 4D). We thus concluded that TEs harboring H3K27me3 contain *cis*-elements bound by transcription factors involved in the regulation of neighboring genes by PRC2.

**Figure 4.**
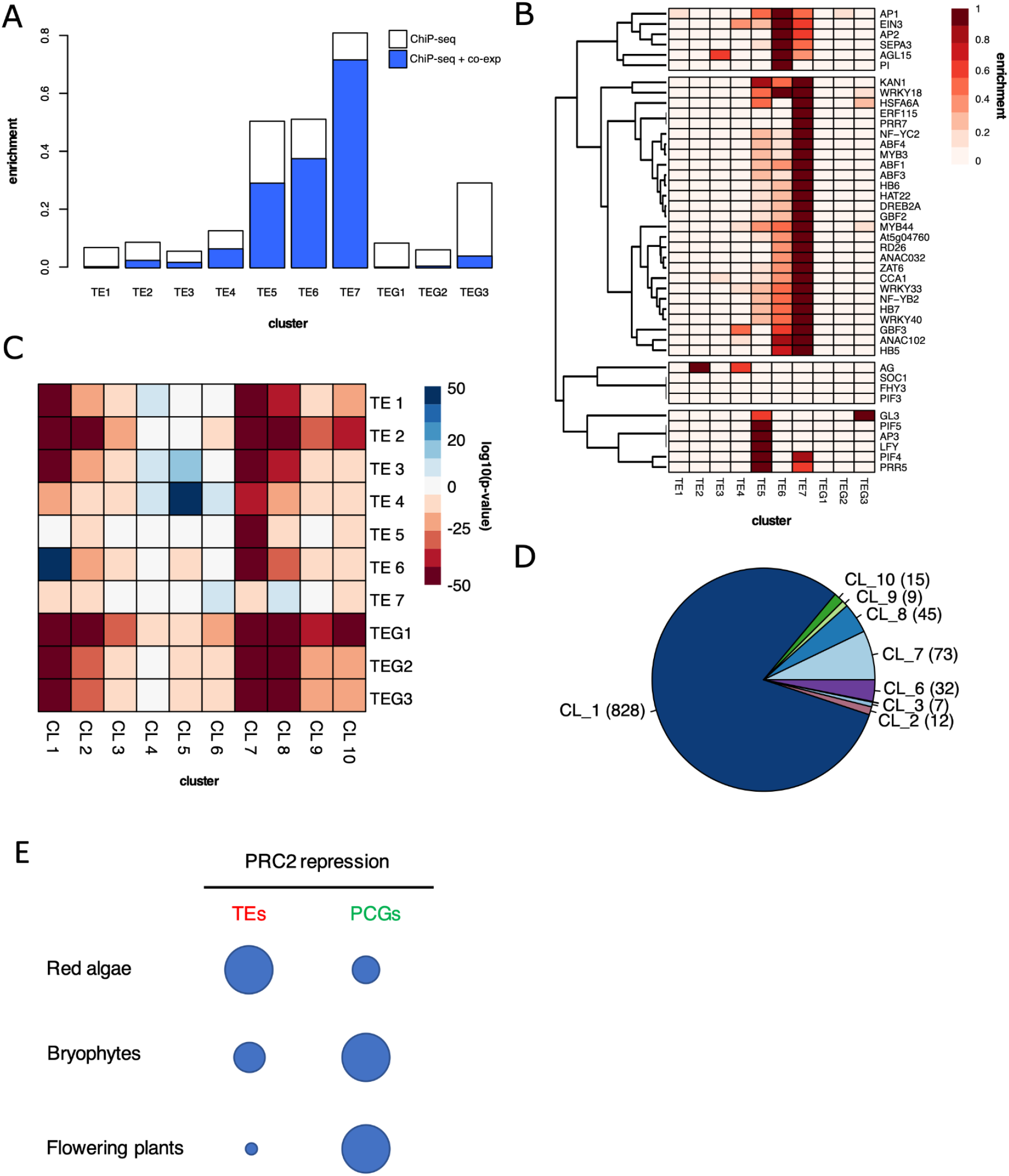
TE fragments are covered by H3K27me3 in Arabidopsis. (A) Bar graph showing enrichment of TF binding sites in each TE and TEG (transposon genes) cluster. White bars indicate TF binding based on all binding events analyzed, blue bars indicate only binding events associated with co-expression of the downstream gene. (B) Heatmap showing enrichment of every TF analyzed in each TE and TEG cluster. (C) Heatmap showing statistical enrichment of every TE cluster in each gene cluster described in ^32^. (D) Pie chart showing proportion of gene clusters of nearby genes of TEs in TE cluster 6 in *Arabidopsis*. (E) A schematic phylogenetic tree illustrating the deep evolutionary conservation of the role of PRC2 in TE silencing with considerable variation among Archaeplastida. Depicted is the number of repressed TEs and PCGs, respectively, in the red alga *C. merolae*, the bryophyte *M. polymorpha*, and the flowering plant *A. thaliana*.

We also noticed a strong enrichment of TF binding in TEs from clusters 5 and 7 (Figure 4A). TEs from cluster 5 are short and marked by an hybrid chromatin state enriched in both H3K9me1 and H3K27me3 (Figures S4A and S4D). They are bound by TFs including the MADS-box factor APETALA3, the bHLH factor GLABRA3 and the pioneer factor LEAFY ^35,36^ that are regulated by H3K27me3 and also control genes regulated by PRC2 and involved in flower development ^33,37^.

The largest group of TFs enriched in TEs belong to the TE cluster 7 (Figure 4A), which are short DNA and RC TEs, not occupied by marks of heterochromatin but with high accessibility at their boundaries (Figure S4). These TEs were strongly associated either with PCGs associated with the Chromatin Landscapes 6 or 8 (Figures 4C and S4E), suggesting that they also contain TF-binding cis-elements controling expression of neighbor PCGs. Overall our results suggest that in Arabidopsis a large number of TFs were domesticated and involved in activation (cluster 7) or PRC2-targeted repression (clusters 5 and 6) of PCGs.

## Conclusions

In summary, we show that PRC2 silences TEs among diverse groups of Archaeplastida. In the red alga *C. merolae*, PRC2 deposits H3K27me3 on a majority of the TEs. Although the proportion of TEs covered by H3K27me3 in bryophytes and angiosperms is less prominent than in *C. merolae*, it is likely that the deposition of H3K27me3 on TEs by PRC2 is general in Archaeplastida. PRC2 also silences TEs in ciliates ^7^ and is associated with TEs in stramenopiles and fungi ^16^ allowing us to propose that PRC2 targeted TEs for silencing in LECA, the common ancestors of these groups. In the ciliate *Paramecium aureliae* the recruitment of PRC2, which deposits both H3K9me3 and H3K27me3 ^7^ involves non-coding RNAs ^8^ in a manner reminiscent of the recruitment of the machinery that deposits H3K9me3 in fungi and animals ^38^. In red algae H3K27me3-silenced TEs are associated with the telomeres which is reminiscent of PRC2 recruitment by TELOMERE REPEAT BINDING FACTORs (TRBs) at telobox motifs in angiosperms ^39^. However we could not find homologs of TRBs in *C. merolae* suggesting that diverse mechanisms of recruitment of PRC2 have been selected to achieve silencing of TEs across eukaryotes.

It has been proposed that during evolution of eukaryotes remnants of TEs became *cis*-elements bound by transcription factors ^40^. Such a scenario has been validated by numerous examples in mammals and accompanied evolution of mammalian specific developmental features such as the placenta and specific aspects of brain development ^1^. In *A. thaliana* a few scattered examples of TE fragments have been associated with specific TF binding sites ^41,42^. Here the presence of TF binding sites on several hundreds of TEs present outside of constitutive heterochromatin support that during plant evolution TEs were domesticated, thus providing an important source of *cis*-elements for TFs that activate or repress transcription.

In plants several examples have illustrated that TEs recruit DNA methylation and H3K9 methylation that affect the expression of nearby PCGs ^40^. These events have been associated with selection of characters in crops in a relatively short period of time ^43–46^. We propose that over very long periods of evolution the recruitment of the ancestral transcriptional repressive Polycomb machinery TEs have been co-opted to silence PCGs. In contrast with the red alga *C. merolae* with TEs enriched in telomeres, in bryophytes, H3K27me3 co-regulates numerous pairs of TEs and PCGs distributed uniformaly across the chromosome, suggesting that TEs contain or behave as *cis*-elements controlling the transcriptional activities of PCGs. In Arabidopsis TEs are bound by transcription factors that recruit PRC2 depositing H3K27me3 on the contiguous PCGs and these transcription factors control expression of genes regulating flower development also controlled by PRC2 ^47^. Interestingly, a recent report showed that a transposon-derived gene controlled by PRC2 was selected as regulator of a reproductive trait in rice ^48^ suggesting that TE-derived cis-elements and proteins coding genes controlled by PRC2 contributed to developmental regulation in plants. These were likely the source of Polycomb gene networks that control multiple aspects of development in land plants and response to environmental cues ^15,21,49^. The ancestral role of PRC2 in TE silencing in distant lineages of eukaryotes suggests that similar evolutionary trajectories might be observed in phyla other than Archaeplastida.

## Acknowledgements

We thank Matt Watson for suggestions and critical reading of the manuscript. F.B. acknowledges support from the PlantS, Next Generation Sequencing and histopathology facilities at the Vienna BioCenter Core Facilities (VBCF), and the BioOptics facility and Molecular Biology Services from the Institute for Molecular Pathology (IMP), members of the Vienna BioCenter (VBC), Austria. This work was funded by FWF grants, P32054 and P33380 to F.B., FWF doctoral school DK W1238 to S.A.M., funding from the European Union’s Framework Programme for Research and Innovation Horizon 2020 (2014‐2020) under the Marie Curie Skłodowska Grant Agreement Nr. 847548 (VIP2) to T.H. and S.W., and funding from the Australian Research Council DP210101423 to J.L.B and a European Research Council, Advanced Grant (project number 787613) from the European Commission to L.D. D.S. acknowledges support by CRC973 (DFG). T.K. was supported by an Elsa-Neumann-fellowship.

## Author contributions

T.H., D.S. and F.B. conceived and designed the experiments. T.F., S.H. and S.M. obtained the *Cme(z)-1* mutant in *C. merolae*. T. K. and R.L. obtained the *Cm(e)z-2* mutant, T.K. performed the RNA-seq of *Cme(z)-1*, R.L. generated western blots in *C. merolae* and *M. polymorpha.* T.H. and S.A.M. obtained the KO mutants in *M. polymorpha*. T.H. performed RNA-seq of *M. polymorpha*. F.R. analyzed the TFs present in TEs in *A. thaliana* marked by H3K27me3 selected by B.J.. T. D. and J. L. B. obtained data that led to the strategy from KO PRC2 in *M. polymorpha*. S. W. annotated TEs in the genomes of *C. merolae*, *A. agrestis* and *M. polymorpha*. T.H. and S. W. performed the analysis of the data with help from E. A. who also performed statistical analyses and curated data. F.B. and D.S. supervised the study. T.H. and F.B. wrote the manuscript draft. The draft was revised with input from D.S., J. L. B. and L. D.

## STAR Methods

### *Annotation of TEs in* C. merolae, A. agrestis *and* M. polymorpha

Transposable elements of *C. merolae* and *M. polymorpha* were annotated using EDTA 1.9.9 ^50^, which incorporates a bunch of tools including LTRharvest, LTR_FINDER, LTR_retriever, Generic Repeat Finder, TIR-Learner, MITE-Hunter, HelitronScanner, and RepeatMasker. All softwares are adjusted to EDTA with proper filters and parameters. Final non-redundant TE libraries are produced by removing nested insertions and protein-coding genes by EDTA customized scripts.

### Plant and alga materials

*Marchantia polymorpha L. subsp. ruderalis* accessions Takaragaike 1 (Tak-1) and Takaragaike 2 (Tak-2; Ishizaki et al., 2016) were used as the wild-type male and female, respectively. Plants were cultured on half-strength Gamborg’s B5 medium solidified with 1% (w/v) agar under continuous white light at 22°C. *Cyanidioschyzon merolae* 10D and Cm*e(z)* cells were grown in liquid culture in sterile modified 2x concentrated Allen’s medium (MA2, ^51^) at 2.5 < pH < 3.0 under constant white light (80 μmol/m^2^/s) at 42°C. Cultures were kept in 50 ml falcon tubes aerated with ambient air supplied through a 1ml serological milk pipette coupled to an aquarium pump. No additional CO_2_ was supplied.

### *Re-analyses of DNA methylation in* C. merolae

Bisulfite-seq data of *C. merolae* were downloaded from the sequence read archive of NCBI under the study PRJNA201680 ^19^. Reads were trimmed with Trim Galore (https://github.com/FelixKrueger/TrimGalore). Bisulfite converted reference genome was prepared from *C. merolae* ASM9120v1 genome sequence using Bismark ^52^. Trimmed reads were mapped to the bisulfite genome using Bowtie2 option of Bismark. Duplicates were removed using deduplicate function in Bismark. Cytosine methylation reports were created from deduplicated reads using bismark_methylation_extractor function in Bismark.

### *Generation of* Cme(z) *mutant*

To inactivate the Cm*E(z)* (CMQ156C) gene, the chromosomal Cm*E(z) orf* was replaced by the *C. merolae URA5.3* selectable marker gene by homologous recombination as follows. All the primers used are listed in Table S1.

To obtain Cm*e(z)-1*, the Cm*E(z)* genomic region, which contained the Cm*E(z) orf* and its 1-kb each of 5’- and 3’-flanking sequences, was amplified with the primers E(z)_KO_F1 and E(z)_KO_R1. The amplified DNA was cloned into the vector pUC19 by In-Fusion Cloning Kit (Takara Bio Inc., Japan). The 5’-flanking sequence of Cm*E(z) orf*, the vector, and the 3’-flanking sequence of Cm*E(z) orf* were amplified with the primers E(z)-KO_F2 and E(z)_KO_R2 and then the *URA5.3* gene, which was amplified with the primers URA_F and URA_R was inserted between the 5’- and 3’-flanking sequence of Cm*E(z) orf* by In-Fusion Cloning Kit. The Cm*E(z)* genomic region, in which Cm*E(z) orf* was replaced with *URA5.3,* was amplified with the primers pUC19_F and pUC19_R and was transformed into *C. merolae* M4, a derivative of *C. merolae* 10D, which has a mutation in the *URA5.3* gene ^53^.

To obtain Cm*e(z)*-2 mutant, 0.5 kb of each, 5′and 3′-sequences, flanking the Cm*E(z) orf* were amplified using the primer sets 5′UTR_E(z)_for/ 5′UTR_E(z)_rev and 3′UTR_E(z)_for/ 3′UTR_E(z)_rev, respectively. The amplified DNA of the 5′ and 3′ flanking region (putative untranslated region, UTR) was cloned into the *SwaI* and *PacI* site, respectively, of the plasmid pSR875 via ligation independent cloning (LIC) method. pSR875 was a kind gift of Stephen Rader and Martha Stark (University of Northern British Columbia, Canada). pSR875 is made from the pBS backbone with *SwaI* and *PacI* LIC sites added + a 10xHis tag + Nos terminator + the Ura5.3 cassette of *C. merolae*. The Cm*E(z)* genomic region, in which Cm*E(z) orf* was replaced with *URA5.3,* was amplified with the primers Cm_trafo_A and Cm_trafo_B was transformed into *C. merolae* T1, a derivative of *C. merolae* 10D, which has a deleted *URA5.3* gene ^54^. Transformation and selection of the gene knockouts were performed as described ^55^.

### *Generation of transcriptome of* C. merolae

*C. merolae* cells were sampled as follows: 2 ml of culture grown to OD_750nm_ = 1 were harvested by centrifugation 3000 x g for 3min at 4°C. The supernatant was removed and the pellet-containing reaction tube put into liquid nitrogen for 15 seconds for cell homogenization purposes. RNA was isolated from frozen pellets using the INNUprep RNA Kit (Analytik Jena) according to the manufacturers’ instructions with the RL buffer supplemented with 10μl/ml beta-mercaptoethanol (β-ME) to improve RNA quality. RNA was eluted in RNAse-free water. Genomic DNA was removed from 1 μg prepared RNA via treatment with DNAse I (Thermo Fisher Scientific) according to manufacturers’ instructions. RiboLock RNase inhibitor (Thermo Fisher Scientific) was added to a final concentration of 1 u/μl. mRNA library was prepared using polyA enrichment and the NEB Ultra RNA Library Prep Kit producing unstranded data. Sequencing was done on an Illumina NovaSeq 6000 platform resulting on 150 bp long paired-end reads. 3G of raw data per sample was produced.

### ChIP-seq data analyses

Details for preparation for ChIP-seq libraries of *A. agrestis* are in ^23^. Chip-seq data of *C. merolae* were downloaded from the Gene expression omnibus of NCBI under the series GSE93913. The bam files of ChIP-seq reads were sorted with SAMtools v1.9 ^56^ and converted to fastq format using bamtofastq function of BEDTools v2.27.1 ^57^, and then trimmed with Cutadapt v1.18 ^58^ and aligned to *A. agrestis* Oxford strain genome ^59^ or *C. merolae 10D* genome (ASM9120v1) ^60,61^ using Bowtie2 v2.3.4.2 ^62^. Resulting bam files were sorted and indexed with SAMtools v1.9. Reads with MAPQ less than ten were removed with Samtools v1.9 and duplicates were removed with Picard v2.18.27 (http://broadinstitute.github.io/picard/). Deduplicated reads from 2 (for *A. agrestis*) or 3 (for *C. merolae*) biological replicates were merged.

H3K27me3 broad peaks of *C. merolae* were called by using macs2 v2.2.5 ^63^. Coverage of H3K27me3 over PCGs and TEs are calculated using the intersect function of BEDtools v2.27.1. PCGs and TEs are considered as covered by H3K27me3 when more than 50% of the regions of each PCG or TE are overlapped by H3K27me3 peaks. Circos plots of PCG annotation, TE annotation and H3K27me3 coverage were made by using circlize package in R ^64^. Chromosome plots were made by using the Chromomap package in R ^65^.

The read coverage of each chromatin mark in *A. agrestis* was normalized against the read coverage of H3 with bamCompare function in deepTools v3.3.1 ^66^, generating bigwig files. K-means clustering of chromatin marks was performed using deepTools v3.3.1. Matrices were computed using computeMatrix for either PCGs or TEs using bigwig files as input and each region is scaled to 2 kb for PCGs and 1 kb for TEs with 1 kb upstream and downstream. Aggregate profile plots of matrices were plotted with plotProfile with k-means clustering. Cluster assignments can be found in ref. Closest function in BEDTools v2.27.1 was used to define the closest TE and gene pair.

### Generation of Marchantia PRC2 knockout mutant

All the primers used to generate *M. polymorpha* PRC2 knockout mutants are listed in Table S1. A DNA fragment producing MpKNOX2-targeting gRNAs was prepared by annealing a pair of synthetic oligonucleotides (TH637/TH638). The fragment was inserted into the BsaI site of pMpGE_En03 (cat. no. 71535, Addgene, Cambridge, MA) to yield pMpGE_En03-MpKNOX2ge1, which was transferred into pMpGE010 (cat. no. 71536, Addgene) (Sugano et al., 2018) using the Gateway LR reaction (Thermo Fisher Scientific, Waltham, MA) to generate pMpGE010_MpKNOX2ge1. This construct was introduced into Tak-2 gemmae using the G-AgarTrap method ^67^. Transformants were selected on 0.5 Gamborg B5 plates without vitamins (Duchefa Biochemie) supplemented with hygromycin and genotyped using the following primer pair: TH652/TH653, leading to isolation of Mp*knox2-1* allele.

To construct a plasmid to disrupt MpE(z)1, two DNA fragments producing MpE(z)1-targeting gRNAs were prepared by annealing pairs of synthetic oligonucleotides (MpEz1-gRNA-1-Fw/MpEz1-gRNA-1-Rv and MpEz1-gRNA-4-Fw/ MpEz1-gRNA-4-Rv). The fragments were inserted into the BsaI sites of pMpGE_En04 and pBC-GE14 to yield pMpGE_En04-MpEz1ge1 and pBC-GE14-MpEz1ge4, respectively. These two plasmids were assembled via BglI restriction sites and ligated to yield pMpGE_En04-MpEz1-ge1-ge4. The resulting DNA fragment containing two MpU6promoter-gRNA cassettes was transferred into pMpGE011 (cat. no. 71536, Addgene) using the Gateway LR reaction (Thermo Fisher Scientific) to yield pMpGE011_MpEz1-ge1-ge4. This construct was introduced into Mp*knox2-1* gemmae using the G-AgarTrap method. Transformants were selected for on 0.5 Gamborg B5 plates without vitamins (Duchefa Biochemie) supplemented with chlorsulfuron and genotyped using the following primer pair: TH650/TH651, leading to isolation of Mp*knox2-1* Mp*e(z)1-1* and Mp*knox2-1*Mp*e(z)1-2* alleles.

To construct a plasmid to disrupt MpKNOX2 and MpE(z)1 simultaneously, a DNA fragment producing MpKNOX2-targeting gRNAs was prepared by annealing a pair of synthetic oligonucleotides (TH637/TH638). The fragment was inserted into the BsaI site of pBC-GE14 to yield pBC-GE14-MpKNOX2ge1. This plasmid was assembled with pMpGE_En04-MpEz1ge1 via BglI restriction sites and ligated to yield pMpGE_En04-MpEz1-ge1-MpKNOX2ge1. The resulting DNA fragment containing two MpU6promoter-gRNA cassettes was transferred into pMpGE010 (cat. no. 71536, Addgene) using the Gateway LR reaction (Thermo Fisher Scientific) to yield pMpGE010_MpEz1ge1-MpKNOX2ge1. This construct was introduced into Tak-1 gemmae using the G-AgarTrap method. Transformants were selected for on 0.5 Gamborg B5 plates without vitamins (Duchefa Biochemie) supplemented with hygromycin and genotyped using the following primer pairs: TH650/TH651 for MpE(z)1, TH652/TH653 for MpKNOX2, leading to isolation of Mp*knox2-2* Mp*e(z)1-3*.

### *Generation of transcriptome of* M. polymorpha *PRC2 mutants*

14 day old plants grown from gemmae were collected and frozen in liquid nitrogen in Precellys tubes (Bertin Instruments, Montigny-le-Bretonneux, France) with 2.8mm zirconium oxide beads (Bertin Corp., Rockville, MD, USA) and disrupted with a Precellys Evolution tissue homogenizer (Bertin Instruments) using the following settings: 4500 RPM 30s, 5s pause, repeated twice. Total RNA was extracted using a Spectrum Plant Total RNA kit (Sigma Aldrich, Merck KGaA, Darmstadt, Germany). Extracted RNA was treated by DNA-free™ DNA Removal Kit (Thermo Fisher Scientific). RNA-seq libraries were generated from 1 μg of total RNA using NEBNext® Ultra™ II Directional RNA Library Prep Kit for Illumina® (New England Biolabs). These libraries were sequenced on an Illumina NextSeq 550 to generate 75bp paired-end reads. Three biological replicates each of Mp*knox2-1* and Mp*knox2-1* Mp*e(z)1-1* were used for subsequent analyses.

### Transcriptome data analysis

Bam files of RNA-seq reads were sorted with SAMtools v1.9 ^56^ and converted to fastq format using bamtofastq function of BEDTools v2.27.1 ^57^, and then trimmed with Trim Galore (https://github.com/FelixKrueger/TrimGalore) and aligned to MpTak1v5.1r2 genome ^9^ for *M. polymorpha* or *C. merolae* ASM9120v1 genome for *C. merolae* using STAR ^68^. Reads counts for genes were calculated by using RSEM ^69^ and those for TEs were calculated by using TElocal (https://github.com/mhammell-laboratory/TElocal). Calculated read counts were imported into R v3.5.1 (R Core Team, 2018) and Differential gene analysis was performed using DeSeq2 v1.22.2 ^70^.

### Real time RT-PCR

Total RNAs were prepared from 14 day old plants grown from gemmae of Tak-2, Mp*knox2-1* and Mp*knox2-1* Mp*e(z)1-2* following the protocol described above. Extracted RNA was treated by DNA-free™ DNA Removal Kit (Thermo Fisher Scientific). cDNA was synthesized from 0.5 μg of total RNA using RevertAid H Minus First Strand cDNA Synthesis Kit (Thermo Fisher Scientific). Real-time RT-PCR analysis was performed using LightCycler® 96 (Roche) and Luna® Universal qPCR Master Mix (New England Biolabs) with the primers listed in Table S1. The analysis was done with three technical replicates and three biological replicates for each genotype. Expression levels of each TEs were normalized against the expression level of MpEF1α. Expression levels of each TE in each genotype were then normalized with the expression level in Mp*knox2-1* Mp*e(z)1-2* and plotted using the ggplot2 ^71^ package in R.

### Nuclei isolation from Marchantia

To isolate nuclei from vegetative tissue of Marchantia, a method described in a previous study was used with some modifications (ref). 500 mg of thallus tissue from 14 day old plants grown from gemmae was collected in a 15 ml plastic tube and frozen with 6 mm zirconium beads. Frozen tissue was disrupted by vortex (max speed, 30 second repeat 6 times). Disrupted tissue was mixed with 5 ml of lysis buffer (20 mM Tris-HCl pH8.0, 25%[v/v] glycerol, 10 mM MgCl_2_, 250mM sucrose, 5 mM DTT, 1x cOmplete™) using vortex and then filtered through double-layered Miracloth. The flow-through was centrifuged at 1500g for 10 min at 4°C. The supernatant was discarded and the pellet was washed three times in 5 mL of wash buffer (20 mM Tris-HCl pH8.0, 25%[v/v] glycerol, 10 mM MgCl_2_, 0.2% [v/v] TritonX-100, 5 mM DTT, 1x cOmplete™). The final pellet was resuspended in 200 μl 1x Laemmlie buffer in 0.2x PBS and boiled at 95°C for 5min.

### Protein extraction from C. merolae and immunoblot analyses

Algae were grown to OD750=1 and concentrated/diluted accordingly. After centrifugation for 5 min at 10000 g, the pellet was dissolved in 100 μl of 4 M Urea, 100 μl of 2xSDS Loading buffer was added and proteins were denatured at 95 °C for 10 min. After separation of proteins on 15 % SDS-gels, they were blotted Immobilon-FL PVDF membranes (IPFL00010, Merck Millipore). After transfer and activation the membrane was blocked with LiCor Odyssey Blocking Solution. Histone modification specific antibodies (α-H3K27me3, Diagenode C15410195, Lot A0821D, 1:2000; α-H3K9me1, Abcam 8896, Lot GR34167862, 3 μg; α-H3K27me1, Merck Millipore 07-448, Lot DAM1661077, 2 μg) were incubated in PBS Odyssey Blocking solution at 4°C overnight with the membrane, then anti-H3pan (Diagenode, C15200011, Lot 002, 1:2000) antibodies were added for 1h at room temperature. Membranes were washed and incubated with the two secondary antibodies (IRDye 680 RD, LiCor926-68070, 1:150000; IRDye 800 CW, Li-Cor 926-32211, 1:75000) for 1 h at room temperature. After drying of membrane, fluorescent signals were detected in a LiCor Odyssey XF at 700 nm and 800 nm, and quantified with LiCor Empiria Studio Software. For peptide competition, 10 μg peptides (H3K9me1, Diagenode C16000065; H3K27me1, Diagenode C16000045) were incubated with histone modification specific antibody in LiCor Odyssey blocking solution for 30 min at room temperature. Subsequently, the membrane was added and incubated overnight at 4°C.

### TF analysis in Arabidopsis

Transcription factor peak BED files were obtained from ChIP-seq datasets compiled previously ^72–76^. Each significant peak region in the genome was classified according to TE clusters ^32^. Only the center of the peak was considered to classify chromatin states. Enrichment for TF binding was performed as in ^77^ but replacing chromatin states for TEs from each cluster. Briefly, for each TE cluster (te) and TF the following formula was applied: (a_te_/b)/(c_te_/d), where a_te_ is the total number of bases of TF peaks for a given te; b is the total number of bases of peaks for a TF; c_te_ is the total number of bases of the te; d is the total number of bases of all TE clusters.

For co-expressed genes, we first annotated peaks over TAIR10 genome using ChIPseeker ^78^ package in R and extracted the ID of downstream genes and used ATTEDII ^79^ coexpressed gene tables (version Ath-r.c3-1) and included the top 10% of both positive and negatively co-expressed genes. For enrichment of TE clusters in PCG clusters, gene clusters from ^32^ were used. We calculated Fisher’s Exact Test comparing enrichment of gene IDs in annotated TEs against PCG clusters, both for the alternative hypothesis of being greater or less than expected were calculated. The −log10(p-value) was assigned if the alternative hypothesis of being greater has the lowest p-value, or the log10(p-value) if not.

