## Supplemental Figures and Table for "Transposons repressed by H3K27me3 were co-opted as cis-regulatory elements of H3K27me3 controlled protein coding genes during evolution of plants"

**Figure S1. Annotation of TEs and re-analysis of 5mC presence in *C. merolae***

**Figure S2. Short TEs in *A. agrestis* and *M. polymorpha***

**Figure S3 Generation of PRC2 knock out mutant in *M. polymorpha***

**Figure S4 TE clusters in *A. thaliana***

**Table S1. oligos used in this study**

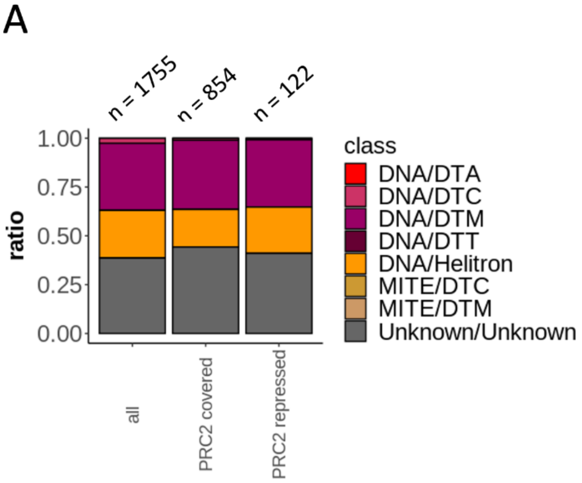


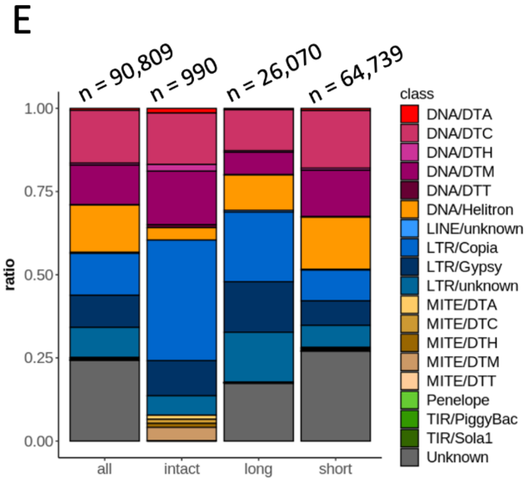


B

| | mCG (%) | mCHG (%) | mCHH (%) |
|---|---|---|---|
| Nuclear | 0.2 | 0.18 | 0.19 |
| Chloroplast | 0.2 | 0.19 | 0.21 |
| Mitochondrion | 0.18 | 0.15 | 0.2 |

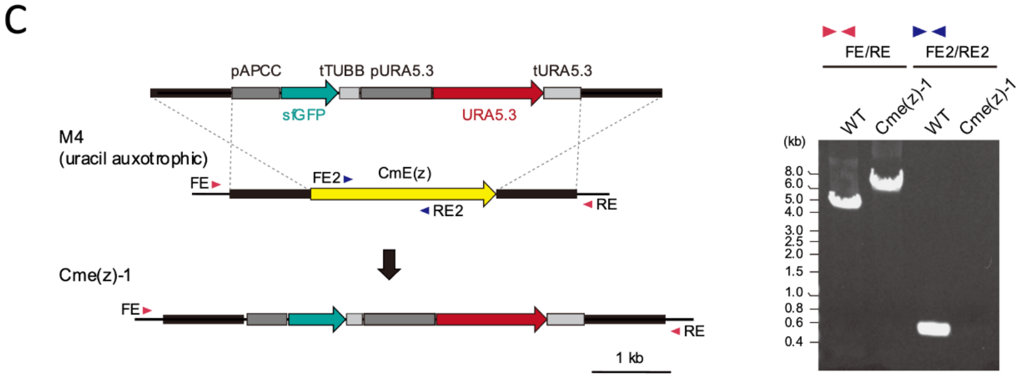


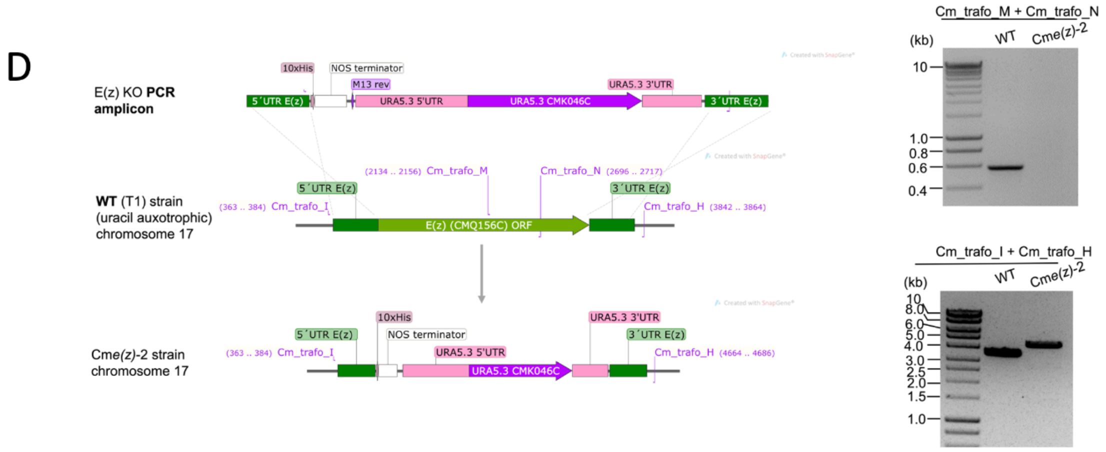

**Figure S1. Annotation of TEs and re-analysis of 5mC presence in *C. merolae***

(A) Stacked bar chart indicating proportion of TE families in all TE annotated (all TE), TEs covered by H3K27me3 (PRC2 covered), or TEs repressed by PRC2 (PRC2 repressed). Total numbers of TEs in each category are shown on bars.

(B) Table showing summary of DNA methylation levels in nuclear and organellar genome of *C. merolae*.

(C), (D) Schematic illustration showing details of Cm*e(z)-1* (C) and Cm*e(z)-2* (D) mutations. Two uracil-auxotrophic WT strains, M4 and T1, respectively, were transformed via homologous recombination resulting in two independent Cm*e(z)* mutant lines.

(E) Stacked bar chart indicating proportion of TE families in all TE annotated (all), intact TEs (intact), TEs longer than 500 bp (long) and TEs shorter than 500 bp (short). Total numbers of TEs in each category are shown on bars.

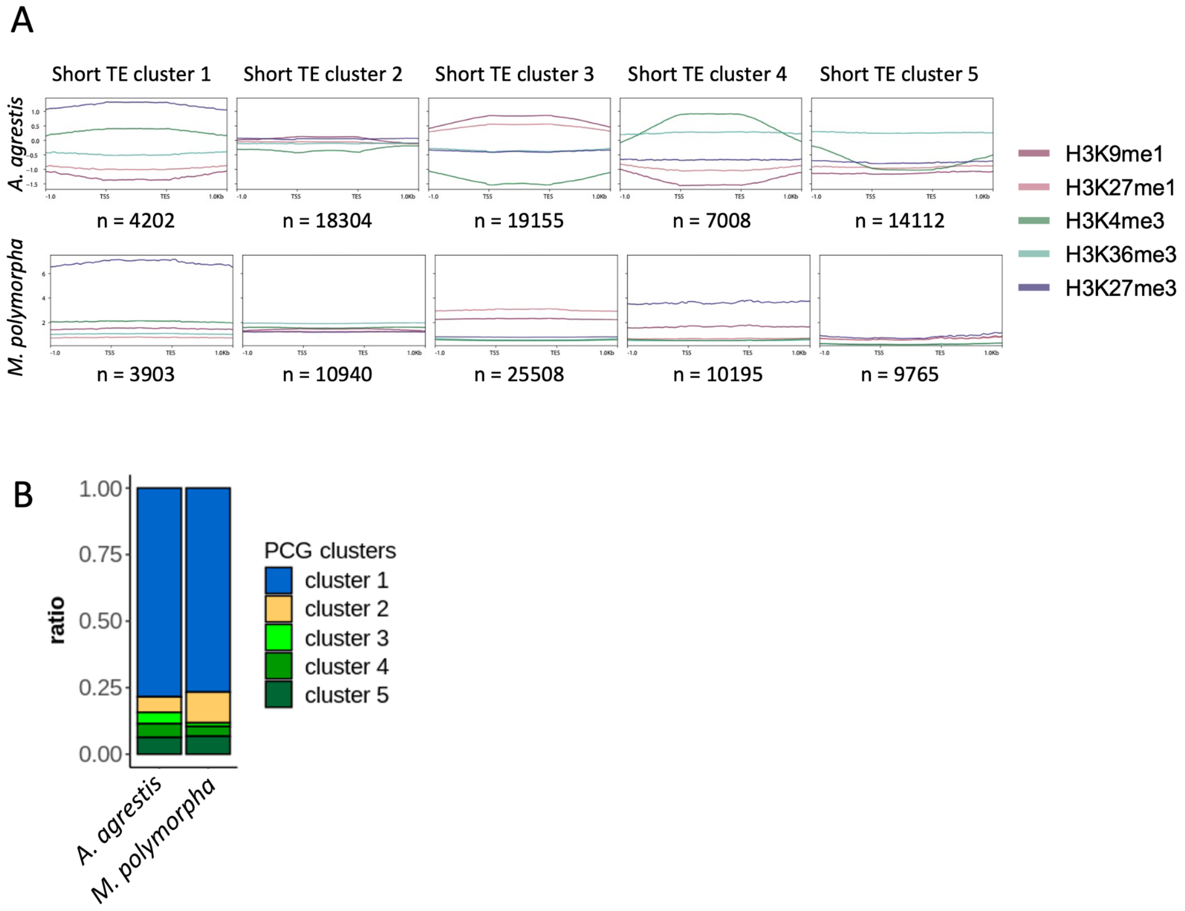


**Figure S2. Short TEs in *A. agrestis* and *M. polymorpha***

(A) Aggregate profile plots showing log2 ChIP/H3 enrichment for various chromatin modifications per short TE clusters of *A. agrestis* (top) or *M. polymorpha* (bottom).

(B) Stacked bar chart showing proportion of gene clusters of nearby genes of TEs in short TE cluster 1 in *A. agrestis* (left) and *M. polymorpha* (right).

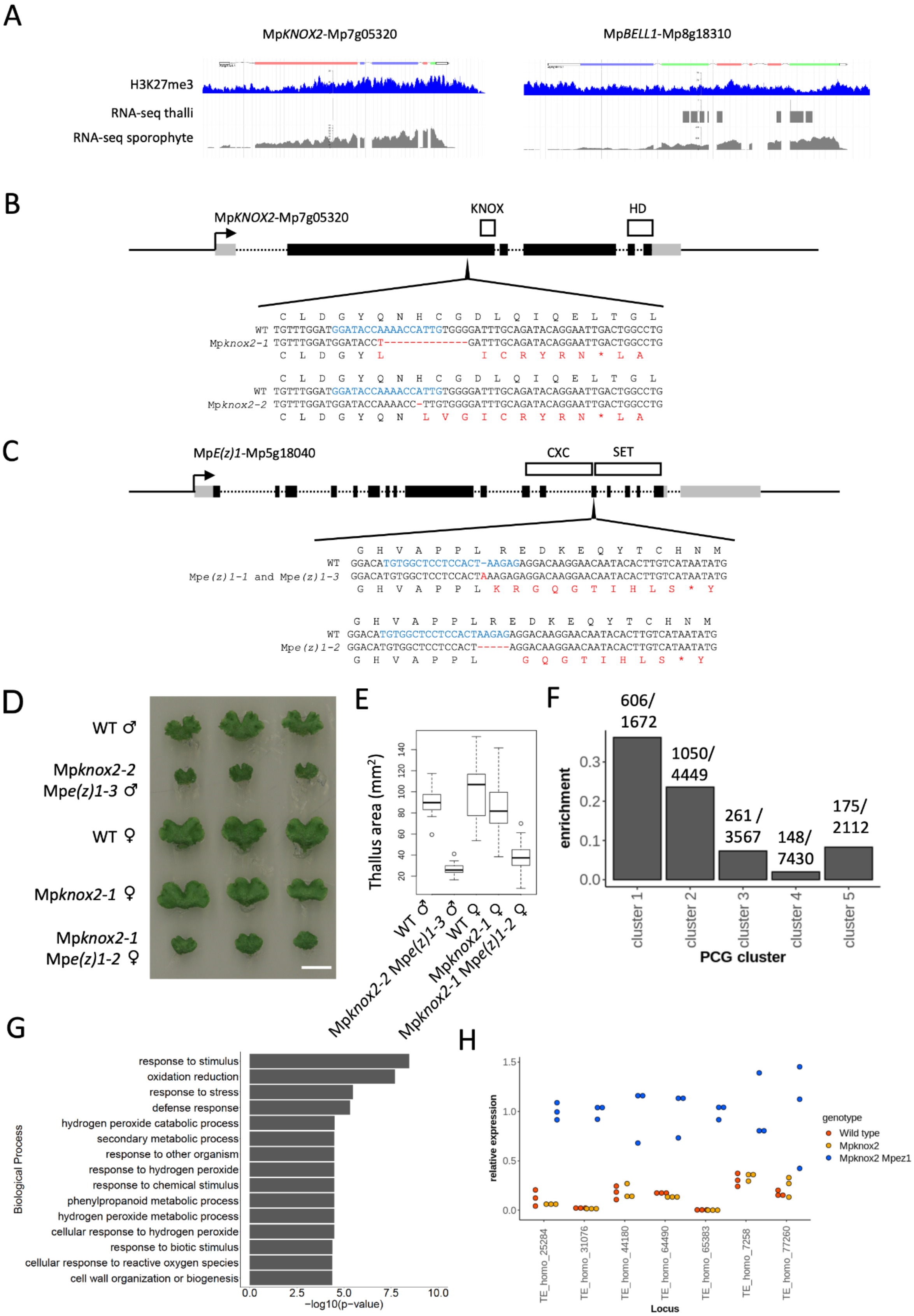

A
MpKNOX2-Mp7g05320
MpBELL1-Mp8g18310
H3K27me3
RNA-seq thalli
RNA-seq sporophyte
B
MpKNOX2-Mp7g05320
KNOX
HD
C L D G Y Q N H C G D L Q I Q E L T G L
WT TGTTTGGATGGATACCAAAACCATTGTGGGGATTTGCAGATACAGGAATTGACTGGCCTG
Mpknox2-1 TGTTTGGATGGATACCT------------GATTTGCAGATACAGGAATTGACTGGCCTG
C L D G Y L I C R Y R N * L A
C L D G Y Q N H C G D L Q I Q E L T G L
WT TGTTTGGATGGATACCAAAACCATTGTGGGGATTTGCAGATACAGGAATTGACTGGCCTG
Mpknox2-2 TGTTTGGATGGATACCAAAACC-TTGTGGGGATTTGCAGATACAGGAATTGACTGGCCTG
C L D G Y Q N L V G I C R Y R N * L A
C
MpE(z)1-Mp5g18040
CXC
SET
G H V A P P L R E D K E Q Y T C H N M
WT GGACATGTGGCTCCTCCACT-AAGAGAGGACAAGGAACAATACACTTGTCATAATATG
Mpe(z)1-1 and Mpe(z)1-3 GGACATGTGGCTCCTCCACTAAAGAGAGGACAAGGAACAATACACTTGTCATAATATG
G H V A P P L K R G Q G T I H L S * Y
G H V A P P L R E D K E Q Y T C H N M
WT GGACATGTGGCTCCTCCACTAAGAGAGGACAAGGAACAATACACTTGTCATAATATG
Mpe(z)1-2 GGACATGTGGCTCCTCCACT-----AGGACAAGGAACAATACACTTGTCATAATATG
G H V A P P L G Q G T I H L S * Y
D
WT ♂
Mpknox2-2 Mpe(z)1-3 ♂
WT ♀
Mpknox2-1 ♀
Mpknox2-1 Mpe(z)1-2 ♀
E
Thallus area (mm²)
20 40 60 80 100 120 140
WT ♂
Mpknox2-2 Mpe(z)1-3 ♂
WT ♀
Mpknox2-1 ♀
Mpknox2-1 Mpe(z)1-2 ♀
F
enrichment
0.0 0.1 0.2 0.3
606/ 1672
1050/ 4449
261 / 3567
148/ 7430
175/ 2112
cluster 1
cluster 2
cluster 3
cluster 4
cluster 5
PCG cluster
G
Biological Process
response to stimulus
oxidation reduction
response to stress
defense response
hydrogen peroxide catabolic process
secondary metabolic process
response to other organism
response to hydrogen peroxide
response to chemical stimulus
phenylpropanoid metabolic process
hydrogen peroxide metabolic process
cellular response to hydrogen peroxide
response to biotic stimulus
cellular response to reactive oxygen species
cell wall organization or biogenesis
0.0 2.5 5.0 7.5 10.0
−log10(p−value)
H
relative expression
0.0 0.5 1.0 1.5
TE_homo_25284
TE_homo_31076
TE_homo_44180
TE_homo_64490
TE_homo_65383
TE_homo_7258
TE_homo_77260
Locus
genotype
Wild type
Mpknox2
Mpknox2 Mpez1

**Figure S3 Generation of PRC2 knock out mutant in *M. polymorpha***

(A) Screenshots of genome browser showing H3K27me3 coverage in thalli (blue) and expression levels in thalli and sporophyte (grey) of Mp*KNOX2* (left) and Mp*BELL1*.

(B), (C) Gene structure of Mp*KNOX2* (B) and Mp*E(z)1* (C). Black lines, grey boxes, black boxes and dotted lines indicate intergenic regions, UTRs, exons and introns, respectively. Regions coding conserved motifs and domains are indicated by white boxes. In sequence alignment, guide RNA target sites are shown in blue and mismatches between wild type and mutants are shown in red.

(D) Images of 14 day old vegetative thalli in each genotype. Three independent plants are shown.

(E) Boxplot showing thallus area of each genotype. n>30

(F) Bar plot indicating enrichment of up-regulated PCGs in each PCG cluster, calculated as the number of PCGs exhibiting increased expression in each cluster divided by the total number of PCGs in each cluster.

(G) GOterm enrichment analysis using PCGs directly repressed by PRC2. Top 15 biological processes are indicated.

(H) Real time RT-PCR analysis confirming increased expression of several intact TE loci in Mp*knox2* Mp*e(z)1*. Expression levels of each TE are normalized against the expression level of the Mp*EF1* gene. In each TE, expression levels are normalized using the expression level in the Mp*knox2* Mp*e(z)1* mutant. Measurements of three biological replicates are plotted.

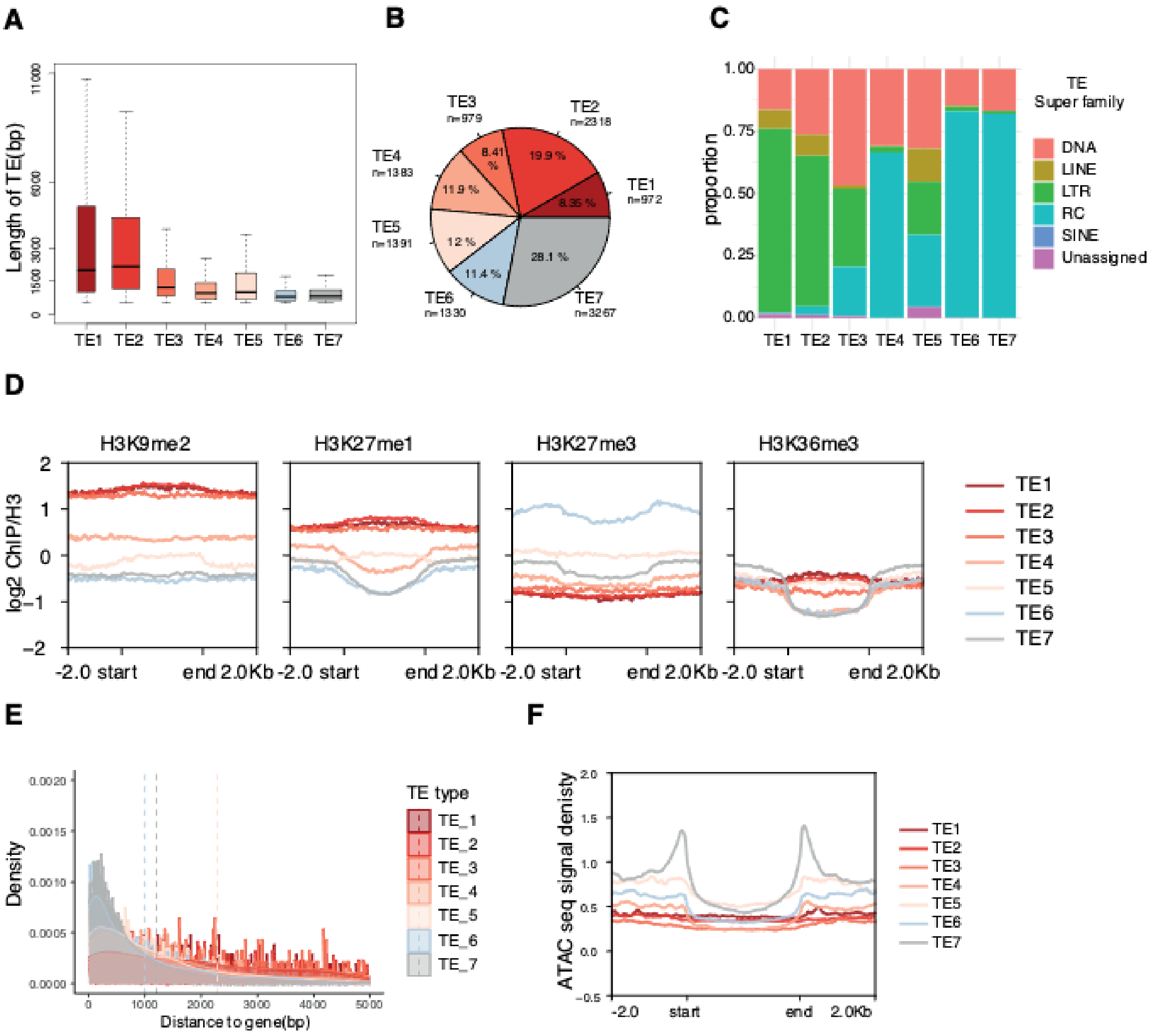


**Figure S4 TE clusters in *A. thaliana***

(A) Boxplot showing length distribution across different groups of TEs defined in [32].

(B) Pieplot showing the genomic abundance of each group of TEs (TE>500 bp).

(C) Barplot comparing the proportions of TEs within each group of TEs w.r.t TE super families.

(D) Aggregate profile plots showing log2 ChIP/H3 enrichment for various chromatin modifications on each group of TEs.

(E) Density plot and histogram showing the distance between TEs and their closest PCG.

(F) Accessibility for each group of TEs.

**Table S1. oligos used in this study**

| name | seq5->3 | description |
|---|---|---|
| E(z)_KO_F1 | cggtacccggggatcCGTTCACCCGACATATGGTCG | To generate Cmez-1 mutant |
| E(z)_KO_R1 | cgactctagaggatcCTCGCTGTTCTATTCGATAGAGG | To generate Cmez-1 mutant |
| E(z)-KO_F2 | agtcagctgctagggCAAGATCCGTGAAGAGAGACGC | To generate Cmez-1 mutant |
| E(z)_KO_R2 | ttcgcccctcagttcATCAATTTTGTGGGGCGCTTTTC | To generate Cmez-1 mutant |
| URA_F | GAACTGAGGGGCGAACGC | To generate Cmez-1 mutant |
| URA_R | CCCTAGCAGCTGACTGTATCTC | To generate Cmez-1 mutant |
| pUC19_F | GCTGCAAGGCGATTAAGTTGGGTAACGCCAGGGTTTTCCC | To generate Cmez-1 mutant |
| pUC19_R | TTATGCTTCCGGCTCGTATGTTGTGTGGAATTGTGAGCGG | To generate Cmez-1 mutant |
| 5´UTR_E(z)_for | CTTATCTCAATATTTGGCCATCGTTTCGGTATCTT | To generate Cmez-2 mutant |
| 5´UTR_E(z)_rev | AACAATCACCAATTTGATCAATTTTGTGGGGCGCTT | To generate Cmez-2 mutant |
| 3´UTR_E(z)_for | AGTTGAAGTATGTTACAAGATCCGTGAAGAGAGACG | To generate Cmez-2 mutant |
| 3´UTR_E(z)_rev | ATGTTAAGTGGATTACGCAAAGAAAGGAGAAAAATGGAA | To generate Cmez-2 mutant |
| Cm_trafo_H | TCTTCGTCGTCGTCAATCCAGCG | To generate Cmez-2 mutant |
| Cm_trafo_I | TCGCAGAGACTCGTTGGGGAAT | To generate Cmez-2 mutant |
| Cm_trafo_M | TTTGTCGTATCCGGCAAGTTTGC | To generate Cmez-2 mutant |
| Cm_trafo_N | CGCAAAGCAGGGACATTTCTTG | To generate Cmez-2 mutant |
| Cm_trafo_A | ggccatcgtttcggtatctt | To generate Cmez-2 mutant |
| Cm_trafo_B | gcaaagaaaggagaaaaatggaatcttgg | To generate Cmez-2 mutant |
| TH637 | ctcgGGATACCAAAACCATTG | gRNA targetting MpKNOX2 |
| TH638 | aaacCAATGGTTTTGGTATCC | gRNA targetting MpKNOX2 |
| MpEz1-gRNA-1-Fw | ctcgTGTGGCTCCTCCACTAAGAG | gRNA targetting MpEz(1) |
| MpEz1-gRNA-1-Rv | aaacCTCTTAGTGGAGGAGCCACA | gRNA targetting MpEz(1) |
| MpEz1-gRNA-4-Fw | ctcgGTAAACAAACATGACTATCT | gRNA targetting MpEz(1) |
| MpEz1-gRNA-4-Rv | aaacAGATAGTCATGTTTGTTTAC | gRNA targetting MpEz(1) |
| TH650 | CCTACGACTGCACCTCCCGAGT | for Mpez1 genotyping |
| TH651 | AAAACGCACCCCACCCAGCTAC | for Mpez1 genotyping |
| TH652 | GTGCAGCAAGTGGAGGCAGGAG | for Mpknox2 genotyping |
| TH653 | CATCAGGCTGGGGGCATTTGGG | for Mpknox2 genotyping |

| | | |
|---|---|---|
| TH749 | CTCTTCGTCCAGTGGGAGGC | TE_homo_64490-Fw |
| TH750 | AGTTCTGGCAGTATTTAAGGAAGGG | TE_homo_64490-Rv |
| TH751 | GTGAGCCGACAGAGTTGAACTCTCC | TE_homo_77260-Fw |
| TH752 | CACTTGGAACGCAAGAGGATTGTGC | TE_homo_77260-Rv |
| TH761 | CCGCTGAAGGGTGAGACAAACAAGG | TE_homo_25284-Fw |
| TH762 | CTCTAGCACCCGAAACTCTGTGTCC | TE_homo_25284-Rv |
| TH696 | CTCCGTGTTGCTTATAAGGTAGCTC | TE_homo_31076-Fw |
| TH697 | CCCATGAGGCACCACTATTAAATTTTG | TE_homo_31076-Rv |
| TH698 | TACAAGCTTGGCCTCGGAATAAATC | TE_homo_44180-Fw |
| TH699 | AGCCATGCATTGTTTGGAAATTTCC | TE_homo_44180-Rv |
| TH708 | GCCTTTCCGTCCCGAGAACAGCTC | TE_homo_65383-Fw |
| TH709 | CGCCATGCACCGAGCTCTGAATTTG | TE_homo_65383-Rv |
| TH702 | GGCTAAGAAACTTCAGGAAGTACTC | TE_homo_7258-Fw |
| TH703 | GCATCGCTGGAAGATGAGCG | TE_homo_7258-Rv |
| MpEF1a-qPCR-Fw | AAGCCGTCGAAAAGAAGGAG | Mp3g23400 |
| MpEF1a-qPCR-Rv | TTCAGGATCGTCCGTTATCC | Mp3g23400 |
